# Chaperone-mediated stress-sensing in *Mycobacterium tuberculosis* enables fast activation and sustained response

**DOI:** 10.1101/2020.06.04.135335

**Authors:** Satyajit D. Rao, Pratik Datta, Maria Laura Gennaro, Oleg A. Igoshin

**Affiliations:** Department of Bioengineering and Center for Theoretical Biological Physics, Rice University, Houston, Texas, USA; Public Health Research Institute, New Jersey Medical School, 225 Warren Street, Newark, NJ 07103, USA

## Abstract

Dynamical properties of gene-regulatory networks are tuned to ensure bacterial survival. In mycobacteria, MprAB-*σ^E^* network responds to the presence of stressors, such as surfactants causing surface stress. Positive feedback loops in this network were previously predicted to cause hysteresis, i.e. different responses to identical stressor levels for pre-stressed and unstressed cells. Here we show that hysteresis does not occur in non-pathogenic *Mycobacterium smegmatis* but occurs in *Mycobacterium tuberculosis*. However, the observed rapid temporal response in *M. tuberculosis* is inconsistent with the model predictions. To reconcile these observations, we implement a recently proposed mechanism for stress-sensing: the release of MprB from the inhibitory complex with chaperone DnaK upon the stress exposure. Using modeling and parameter fitting, we demonstrate that this mechanism can accurately describe the experimental observations. Furthermore, we predict perturbations in DnaK expression that can strongly affect dynamical properties. Experiments with these perturbations agree with model predictions, confirming the role of DnaK in fast and sustained response.

**IMPORTANCE:** Gene-regulatory networks controlling stress response in mycobacterial species have been linked to persistence switches enabling the bacterial dormancy within a host. However, the mechanistic basis of switching and stress sensing is not fully understood. In this paper, combining quantitative experiments and mathematical modeling, we uncover how interactions between two master regulators of stress response -- MprAB two-component system and alternative sigma factor *σ^E^* – shape the dynamical properties of surface-stress network. The result show hysteresis (history dependence) in the response of pathogenic bacteria *M. tuberculosis* to surface stress and lack of the hysteresis in non-pathogenic *M. smegmatis*. Furthermore, to resolve the apparent contradiction between the existence of hysteresis and fast activation of the response, we utilize a recently proposed role of chaperone DnaK in stress-sensing. The results leads to a novel system-level understanding of bacterial stress-response dynamics.

## INTRODUCTION

The intracellular pathogen *Mycobacterium tuberculosis* is highly successful in humans as over a billion individuals are estimated to carry a latent infection. To survive in the host, *M. tuberculosis* must sense stress conditions generated by the host immune system and adapt to them by reprogramming its gene expression and metabolism. Cell-envelope damage is one such stress condition (1, 2). As in many bacteria, the response to this stress involves complex gene regulatory network involving transcriptional master regulators – two-component systems (TCSs) and alternative sigma factors (2, 3, 4).

The core surface-stress response network of *M. tuberculosis* involves the alternative sigma factor *σ^E^* and the MprAB TCS that consists of a histidine kinase (MprB) and a response regulator (MprA). The presence of surface-stressors (surfactants) triggers autophosphorylation of MprB at the histidine residue. The phosphoryl group is then transferred to the Asp residue of MprA. MprB also has phosphatase activity whereby it catalytically dephosphorylates phosphorylated MprA (MprA-P) (5, 6). MprA-P is a transcription factor that activates expression of multiple genes including its own operon, thus creating a positive feedback loop. Another transcriptional target of MprA is *sigE*, gene encoding for *σ^E^*(7). This alternative sigma factor binds core RNA polymerase and guides it to promoters of stress-responsive genes, including the *mprA-mprB* operon. Upregulation of *mprA-mprB* by *σ^E^* results in a second positive feedback loop (4, 8). In addition, *σ^E^* activity is also controlled post-translationally by sequestration by the antisigma factor RseA, which disables *σ^E^* from interacting with RNA polymerase (8, 9).

The MprAB-*σ^E^* stress-response network has recently been linked with persistence, a state in which tubercle bacilli survive inside host immune cells, where they encounter nutrient and oxygen limitation and antibacterial mechanisms (10, 11, 12). The mechanism(s) behind cells switching from bacterial growth to a persistent state remains unknown. However a target of *σ^E^*, relA (a stringent response regulator), showed a bimodal distribution in a population of *Mycobacterium smegmatis* cells (13, 14). A bimodal distribution in agene’s expression level can arise out of bistability in the MprAB-*σ^E^* network (i.e., existence of two distinct states of response for the same level of stress), and a bistable network is a good candidate for a persistence switch if one of the states cease growth. This possibility was explored by previous theoretical study from our team (15). Its results demonstrated that positive feedbacks in this network togetherwith increased effective cooperativity due to RseA-*σ^E^* interaction could result in bistability over a wide range of parameter values. As a result, for a certain range of signals, the transcription activity of *σ^E^* can be in either high (activated) or low (inactivated) depending on initial conditions (see figure 5 in (15)). Bistability would then manifest as hysteresis in response to increasing and decreasing signals, i.e. fully pre-stressed and unstressed cells may show different *σ^E^* activity under identical intermediate stress levels. However, this theoretical prediction of bistability have not been experimentally tested.

Here we investigate whether the predicted hysteresis in the transcription activity of *σ^E^* is observed experimentally in two mycobacterial species, the non-pathogenic *M. smegmatis* and pathogenic *M. tuberculosis*. To investigate the possibility of hysteresis, we examine mycobacterial response to increasing and decreasing surface stress created by different concentrations of surfactant SDS. Furthermore, we compare the model predictions of transient activation or deactivation of *σ^E^* activity following addition or removal of SDS. Using mathematical modeling and parameter fitting, we identify interactions in the network that explain experimentally observed responses. We find that a simple model assuming a first-order activation of MprB autophosphorylation in response to SDS exposure fails to explain the experimental observations. To resolve the inconsistency, we propose a more complex model for MprB activation. This model implements a proposed mechanism involving DnaK, a chaperone that deactivates MprB in unstressed conditions (16). Extra-cytoplasmic proteins unfolded due to surfactant exposure compete with MprB for DnaK, eventually activating MprAB TCS. The model not only explains the observed dynamical properties of the stress response, but also predicts changes in stress response with perturbed DnaK levels. These predictions are confirmed with an engineered strain, confirming the assumptions of the model. Thus, synergistic use of experiments and modeling can uncover interactions in signaling networks shaping their dynamical properties.

## RESULTS

### Lack of hysteresis in transcript levels in *M. smegmatis* can be explained by lack of positive feedback

To experimentally investigate the possibility of bistability, we first examine the response of *M. smegmatis* to increasing and decreasing levels of surfactant SDS that causes cell envelope stress. If our prediction of bistability in this network is correct, we expect to observe hysteresis, i.e., different intermediate states depending on cell history. To test this possibility, previously unstressed or maximally stressed cells were subjected to different intermediate concentrations of SDS. Maximal stress was first identified as the concentration of SDS resulting in bacteriostasis (0.02%, Figure S1). The state of the network was measured as *sigE* transcript abundance following SDS exposure for 2 hours. The results show no hysteresis in *sigE* over the range of SDS concentrations (Figure 1a).

**FIG 1.**
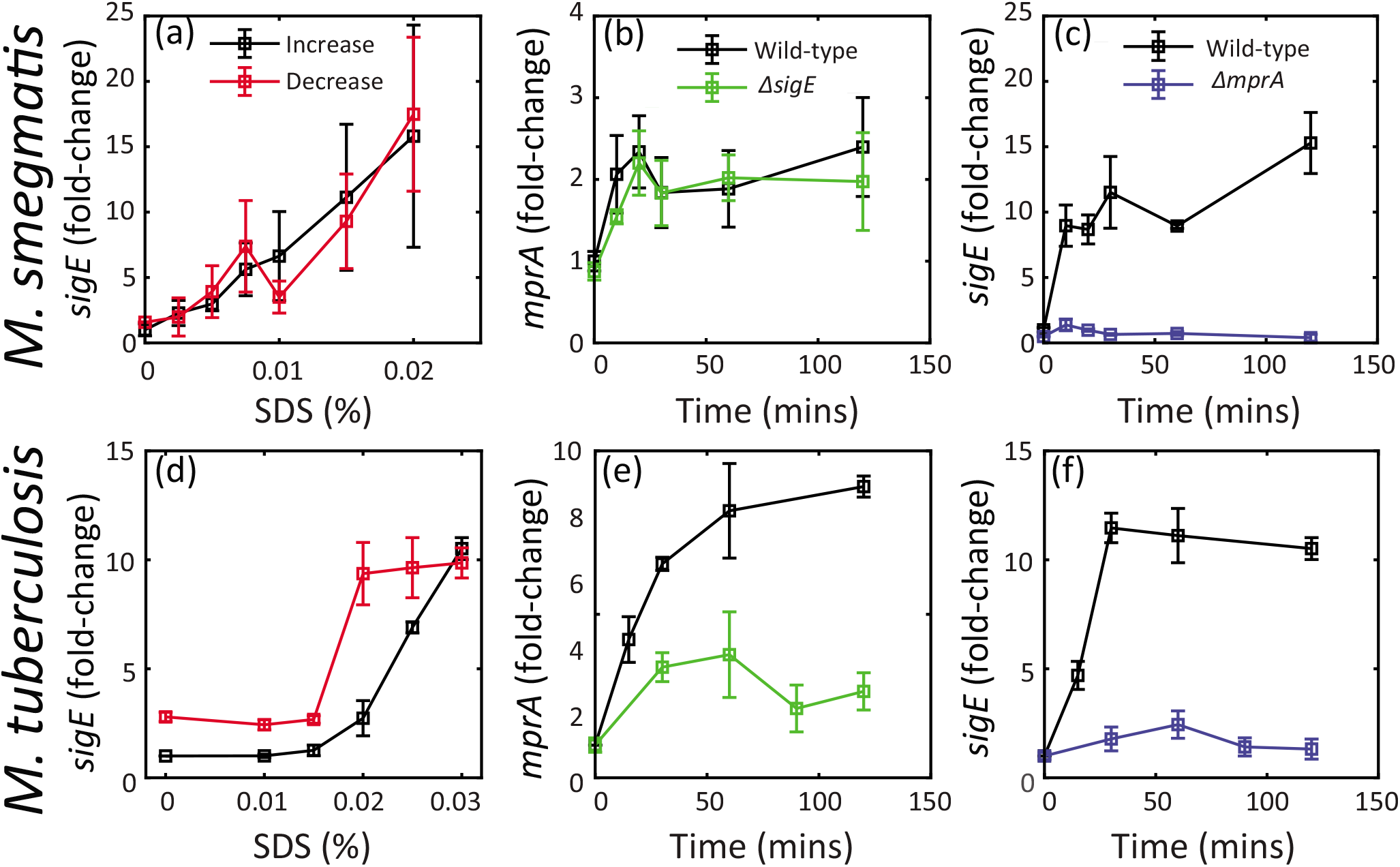
Bacterial transcripts were enumerated by real time PCR, using gene specific probes. Transcripts were normalized to 16S rRNA, and expressed as fold-change relative to pre-treatment. Here and in subsequent figures, mean values (± standard error of the mean) are presented from triplicate experiments. (a) *M. smegmatis* dose-response does not display hysteresis in mRNA levels. Wild-type cells were grown up to mid-log phase and treated with increasing SDS concentrations and harvested 2 hrs post-treatment (black). Wild-type cells were treated with bacteriostatic SDS (0.02%, Figure S1) for 2 hrs, centrifuged, and re-suspended in fresh media containing same or decreasing SDS concentrations and harvested 2 hrs post-treatment (red). (b) Deletion of *sigE* gene does not affect *mprA* time-course, suggesting *M. smegmatis* lacks feedback from *σ^E^* upregulating *mprA*. Mid-log cultures of wild-type and *sigE* deletion mutant were treated with 0.02% SDS and harvested pre-treatment (time 0) and at multiple times post-treatment. (c,f) Similar time course measurement of *sigE* mRNA in wild-type and *mprA* deletion mutant shows very low fold-change in expression suggesting MprA regulates *sigE* expression in both *M. smegmatis* and *M. tuberculosis*. (d)*M. tuberculosis* dose-response displays hysteresis in mRNA levels. (e) Deletion of *sigE* gene affects *mprA* time-course, suggesting *M. tuberculosis* has feedback from *σ^E^* upregulating *mprA*.

Given that positive feedback is required for bistability, we test whether this feedback exists in *M. smegmatis*. We measured *mprA* transcript abundance in wild-type and *sigE*-deletion strains at regular time intervals following exposure to maximal SDS=0.02% (reffig:exptb). At all time points, the transcript abundance of *mprA* in the *sigE* deletion mutant does not deviate significantly from wild type. These measurements indicate that *sigE* deletion has no measurable effect on *mprA* transcription during SDS stress. Additionally, bioinformatics search for a *σ^E^* binding site using consensus sequence described in (4) in the upstream region of *mprA* did not yield any matches. In contrast, *sigE* expression is highly attenuated compared to wild-type in an *mprA* deletion strain (Figure 1c). Altogether, our data show that MprA activates *sigE* transcription but the feedback loop from *σ^E^* to *mprA* is absent in *M. smegmatis*. Thus, lack of bistability, and therefore hysteresis, could be due to a lack of positive feedback.

### Hysteresis observed in *sigE* transcript levels in *M. tuberculosis*

In contrast to the *M. smegmatis* data presented above, reports have suggested that the feedback from *σ^E^* to MprA exists in *M. tuberculosis* (7, 8). Here we confirm these observations demonstrating that transcription of *mprA*, following exposure of *M. tuberculosis* cells to SDS, is significantly decreased in a *sigE* deletion mutant in comparison to wild-type cells (Figure 1e). With the existence of two positive feedback loops confirmed (8, 17), we examined whether hysteresis could be observed in *M. tuberculosis*. We measured *sigE* transcripts in previously unstressed and maximally stressed cells exposed to intermediate concentrations of SDS for 2 hrs. For maximal stress, bacteriostatic SDS at 0.03% was used((18)).The results indicate that, unlike *M. smegmatis*, the *M. tuberculosis* network exhibits hysteresis (Figure 1d). Taken together, our findings in *M. tuberculosis* and *M. smegmatis* strongly suggest a role for positive feedback in hysteresis in mycobacterial response to surface stress. Notably, in pre-stressed cells, *sigE* transcript levels remain above basal levels even after the removal of stressors, i.e. at 0% of SDS (Figure 1d). This contrasts the predictions of our previous model in Tiwari et al. (15). To illustrate this we have simulated dose-response using this model and parameters (Figure 2a). Using MprB autophosphorylation as the signal mimicking the surface stress, we computed the steady state *sigE* mRNA level as a function of signal. Black and red curves correspond to different initial conditions: system can start at the steady state concentrations corresponding to low (black) and high (red) signal respectively (i.e. initially unstressed of maximally stressed cells). The results show that while different steady states are predicted to occur at the intermediate range of signal, *sigE* transcripts return to basal levels when signal is low. This contrasts the observations in Figure 1d. Therefore, the mechanism of hysteresis may be more complicated than the previous model suggested or the model operates in the wrong parameter regime.

**FIG 2.**
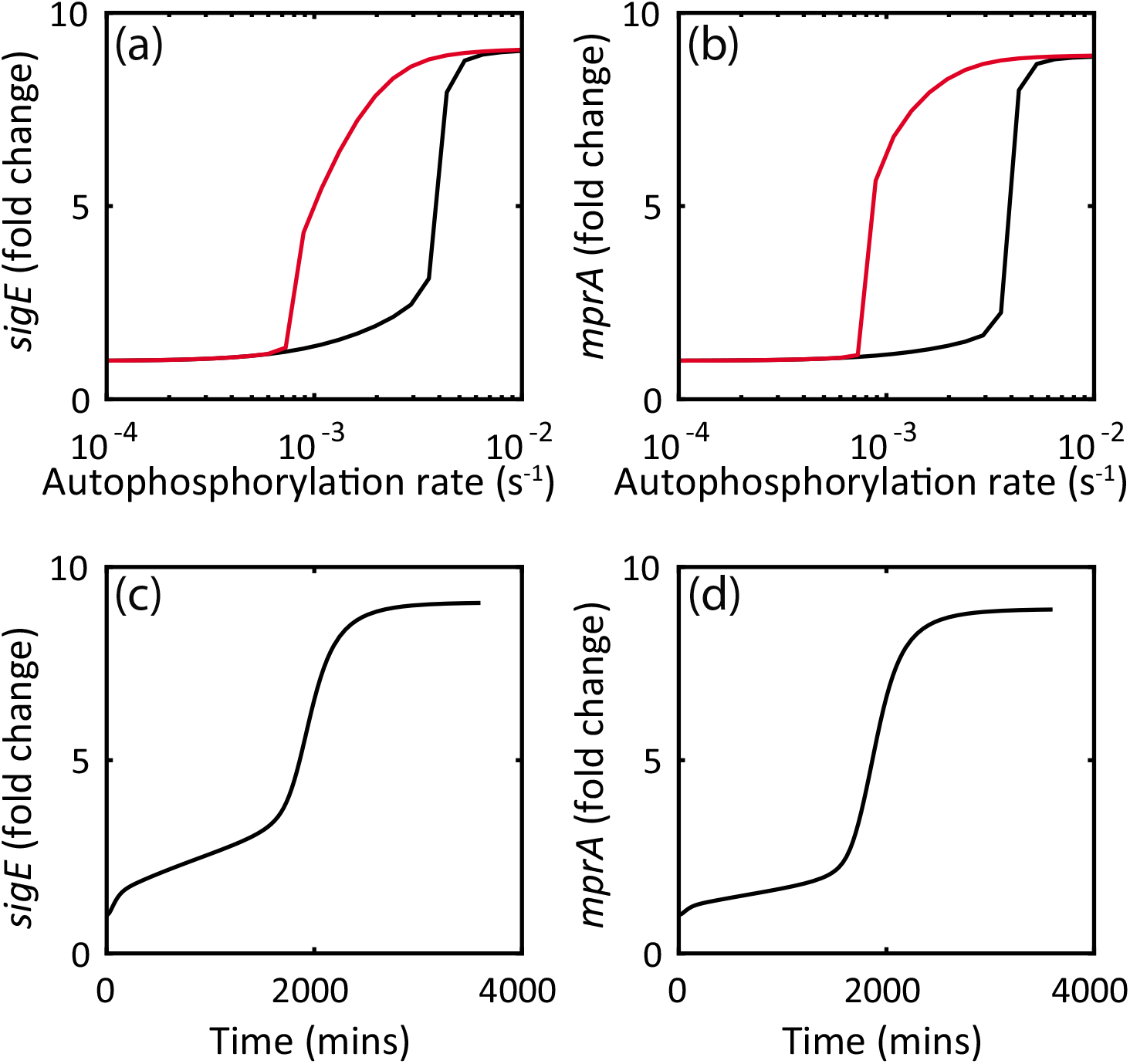
(a,b) steady state dose-response simulations of *mprA, sigE* starting from low (black) and high (red) signal predict hysteresis. (c,d) A very slow response (order of 2000 mins) is predicted for a switch from low to high signal.

### Hysteresis is seen in both *mprA* and *sigE* but the dynamics is unexpectedly fast

To further compare the predictions of our previous model (15) to experimental observations, we test the prediction of hysteresis in *mprA* mRNA (Figure 2b) by measuring *mprA* transcripts in previously unstressed and maximally stressed cells exposed to intermediate SDS concentrations for 2 hrs. The results show that *mprA* transcripts display hysteresis (Figure 3c, triangles). We note that in contrast with *sigE* and in agreement with model prediction, *mprA* transcripts reach basal levels after removing SDS (Figure 3c).

**FIG 3.**
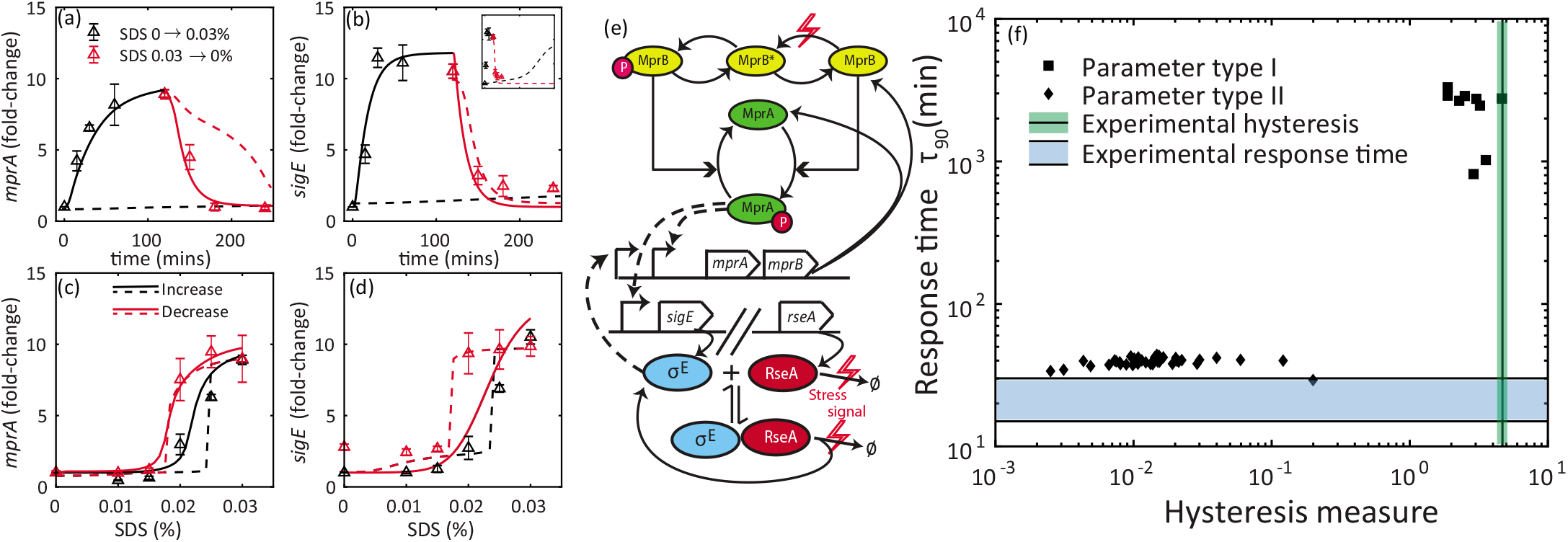
(a-d, triangles) Dose-response and time-course enumeration of transcripts in *M. tuberculosis* (a,b) Time course measurements show rapid accumulation of transcripts. See Experimental Methods for protocols. Cultures were exposed to 0.03% SDS, and *sigE* (MprA-P target) and *mprA* (*σ^E^* & MprA-P target) mRNA levels were enumerated at multiple time points over the course of 2 hours (black triangles). Measurements are presented here as fold-change with respect to pre-treatment levels. Cultures were subsequently re-suspended in fresh SDS-free media to measure a step-down time course (red triangles). (c,d) Dose response measurements show hysteresis in *sigE, mprA* mRNA levels depending on history of stress exposure - previously stressed (red triangles) and unstressed (black triangles). (e) Schematic of the MprAB-*σ^E^* network model. Red lightning symbols represent signal (SDS exposure). SDS exposure leads to MprB switching from phosphatase to kinase form, and also to degradation of RseA (Ø indicates RseA degradation products) and release of *σ^E^*. Dashed arrows indicate transcriptional activation. (a-d, solid lines) Fitting parameters for a simple model of TCS activation (e) results in simulations that match time-course data well (a,b), but dose-response simulation of *sigE* mRNA shows no hysteresis (c,d). (a-d, dashed lines) In models that minimize steady state dose-response (c,d), time course simulation suffers a large activation delay (a,b, dashed black lines). Inset shows activation time on the order of 1000 mins. Simulated mRNA levels were normalized to respective pre-treatment values. (f) Analysis of multiple parameter sets that generate fits of the type in (a-d) illustrates the trade-off between explaining fast response time (diamonds) and hysteresis (squares) in a simple model of MprAB TCS activation shown in (a). Shaded regions - Response time and a measure of hysteresis (see methods) ranges computed from experimental data

In addition to testing model predictions of steady-state responses, we investigated the response dynamics following activating and deactivating signals. Notably, time-course simulation of our previous model(15) shows very slow response (Figure 2c, d). Starting with an initial condition corresponding to low signal (i.e. unstressed condition), we simulate exposure to maximal stress level by step-wise change in MprB autophosphorylation rate to the value corresponding to saturated steady state response (1 s^−1^). The results show unrealistically slow kinetics of mRNA accumulation (Figure 2c,d), with a predicted response time on the order of ~33 hours. If that is true, the protocol used to measure hysteretic response may not be sufficient to achieve steady state. We note that predicted slow response is consistent with expectation of critical slowdown in kinetics around a bistable threshold (19). To experimentally test this prediction, we expose previously unstressed cells to 0.03% SDS and measure *sigE* and *mprA* transcripts at regular intervals for 2 hours following treatment (Methods). The results (Figure 3a,b; black triangles) show that both *mprA* and *sigE* transcripts accumulate quite fast in contrast to model predictions. Thus the discrepancy between observed rate of transcript accumulation and predictions of the our previous model (15) requires us to revisit our network model and parameters.

### Models with simple activation mechanism of MprAB two-component system cannot explain dynamical properties

To understand mechanisms that lead to unexpectedly fast accumulation of target mRNAs, we start with the our previous model (15) with two slight modifications in order to account for two important but previously unaccounted aspects of stress-sensing mechanism(Figure 3e). First, instead of modeling stress by increasing MprB autophosphorylation rate, we assume that stress controls both kinase and phosphatase activity of MprB. In many two-component systems, sensory transduction is driven by a conformation change, enhancing kinase and decreasing phosphatase activity (20, 21, 22). Thus, we include two conformations of MprB explicitly in the model. In the absence of surface stress, MprB is phosphatase-dominant (16) and switches to a kinase form in response to stress. The stress signal modulates first order rate constant of switching between these two forms. Second, we explicitly introduce stress-dependent modulation of RseA activity. In the presence of SDS, RseA has been reported to undergo phosphorylation by a transmembrane kinase PknB and subsequent proteolytic degradation (9). We include this in the model by introducing additional RseA degradation reaction with a signal-dependent rate constant (Methods).

With the revised model, we seek to generate parameters that minimize deviation of simulated time-course and dose-response from experimentally observed data-points. To this end, we employ minimization of the sum-of-square-errors between the model predictions and experimental data-points using particle swarm optimization (Methods). We seek to obtain a large number of parameter sets in this way to account for some parameters having more or less effect on the measured variables. None of the optimized parameters sets is adequate to explain both steady state and dynamic response (Figure 3a-d).

To understand the reasons for this discrepancy, we focus on optimizing these datasets separately. While simulations using optimized parameters can match time-course of mRNA accumulation (Figure 3a,b, solid lines), they lack hysteresis in dose-response, especially in *sigE* (Figure 3 c,d, solid lines). At intermediate SDS, the simulated *sigE* mRNA levels are the same regardless of the initial condition of the network - OFF or ON (Figure 3d, black and red lines overlap). Given that our previous analysis suggested that MprAB-*σ^E^* network can be bistable(15), we attempt to match only dose-response data points at steady state by relaxing the condition for fast mRNA accumulation. Parameter sets that minimize only steady state dose-response error show a close match with experimental data (Figure 3 c,d; dashed lines). However, simulated time-course of mRNA accumulation show a much slower response than the experimental data (Figure 3 a,b; dashed black lines; Figure 3b inset). This suggests that, while the network model can match time-course and dose-response experimental data points separately, it is unable to do so when the two data sets are included simultaneously (discussion for this trade-off follows in the next section). While the simulations in Figure 3 are representative, the trade-off between hysteresis and fast accumulation time is robust. We illustrate this with a scatter plot of a measure for hysteresis and response time ((Figure 3f), Methods). Each point represents a parameter set that fits one data set adequately (either time course (diamonds) or steady-state dose-response (squares)), obtained from runs of particle swarm optimization with random initial seeds. No optimized parameter sets occupy the space at the intersection of experimental hysteresis and response time measures (shaded areas).

### Robustness property of TCSs may lead to response speed-hysteresis trade-off in simple TCS model

To understand why hysteresis is absent when using parameter sets that match time course (solid lines, Figure 3 a-d), we analyzed our ODE model with time-scale separated modules as described previously in (15). We find that the MprAB two-component system lies in a regime of absolute concentration robustness that has been observed in TCSs in bacterial systems (Supporting material, Figure S2)(23, 24, 25). In this regime, the output of a TCS (MprA-P) is invariant to total MprA/MprB concentrations. This in turn ensures that MprA-P output (*sigE* mRNA) is invariant to positive feedback. In contrast, with parameters that describe hysteresis but not fast accumulation (dotted lines, Figure 3 a-d), the TCS lies outside this regime where the MprA-P depends on the total amount of MprA present in cells (Figure S2, dotted lines). In fact, in this regime MprA is saturated and almost all of it is phosphorylated. The modules intersect in 3 points, showing bistability in the network with parameters that display hysteresis (the intermediate intersection represents unstable steady state). The steady state of the network would depend on the initial condition. Thus, we conclude that a fast activating TCS cannot display dose-response hysteresis due to robustness property of two-component systems. Conversely, models displaying hysteresis cannot obtain fast activation dynamics due to being close to bistable threshold (26).

### DnaK-dependent activation of MprB resolves trade-off between response time and hysteresis

To resolve the previously described trade-off, we look for network designs that can generate bistability in a biological network without activation delays. A potential design consists of a transcription factor (TF) sequestered by a stoichiometric inhibitor, coupled with positive autoregulation of TF (19). In absence of activating signal, inhibitor concentration exceeds that of TF keeping it inactive. If activating signals titrate the inhibitors, TF is released and can upregulate its transcriptional target. When positive feedback is present in the network, prolonged exposure to signal can lead to accumulation of TF to the level exceeding these of the inhibitor. In that case, even in the absence of signal there can be a residual TF activity. Furthermore, given that activation is driven by posttranslational sequestration reactions its dynamics should be fast. In fact, such activation mechanisms appear plausible in MprAB-*σ^E^* network. DnaK, a mycobacterial chaperone protein, has been shown to bind to extracytoplasmic domain of MprB and suppress its autokinase activity (16). In an *M. tuberculosis* mutant strain expressing *dnaK* from a chemically-inducible promoter, the MprAB-*σ^E^* network did not activate even after exposure to 0.05% SDS (16), suggesting that the concentration of DnaK is an important factor for stress-response activation. We implement the following mechanism for activation of MprAB based on the results of Bretl et. al (16). MprB can only autophosphorylate when not bound to DnaK and only has phosphatase activity when bound to DnaK. Exposure to SDS increases load of unfolded/misfolded extra-cytoplasmic proteins. Recruitment of chaperone DnaK to those proteins releases MprB to autophosphorylate and activate MprA.

To test whether this modified MprAB TCS can match the experimentally observed hysteresis, we incorporate MprB-DnaK binding in our previous model (Figure 4a). Instead of representing surface stress by a single kinetic rate constant as in the previous model, we model exposure to SDS as a step-increase in a hypothetical DnaK target representing misfolded/unfolded extra-cytoplasmic proteins (see Methods for more details). This increase consequently decreases DnaK concentration available to bind MprB. When implemented, we use this MprAB-DnaK-*σ^E^* model to fit the measured experimental data. As a result, we are able to obtain multiple parameter sets with which the model exhibits hysteresis at intermediate SDS concentrations (Figure 4 b-e) and matches the observed activation kinetics. Bifurcation analysis of the model as a function of signal (SDS concentration) shows that the model is indeed bistable at intermediate signal levels (Figure S3). Notably, when we adapt the MprAB-DnaK-*σ^E^* model to fit *M. smegmatis* data by removing positive feedback from *σ^E^* to MprAB, we can reconcile the absence of hysteresis in dose-response (Figure S4). Using the model and parameters obtained for *M. tuberculosis* however, we find that the models fail to reproduce residual transcriptional activity in pre-stressed cells for small SDS (less than 0.015%) observed in the data (Figure 4e, compare red line with red triangles). Repeating the experiment on 3 different days, we have demonstrated that residual levels of *sigE* mRNA even two hours after complete SDS removal is highly reproducible (Figure 4f). Therefore, we conclude that further modifications of the proposed model are needed to fully explain the data. In our model, only phosphorylated MprA is transcriptionally active. Thus despite accumulation of MprA due to positive feedback, quick dephosphorylation of MprA-P following the SDS removal leads to negligible *sigE* transcription.

**FIG 4.**
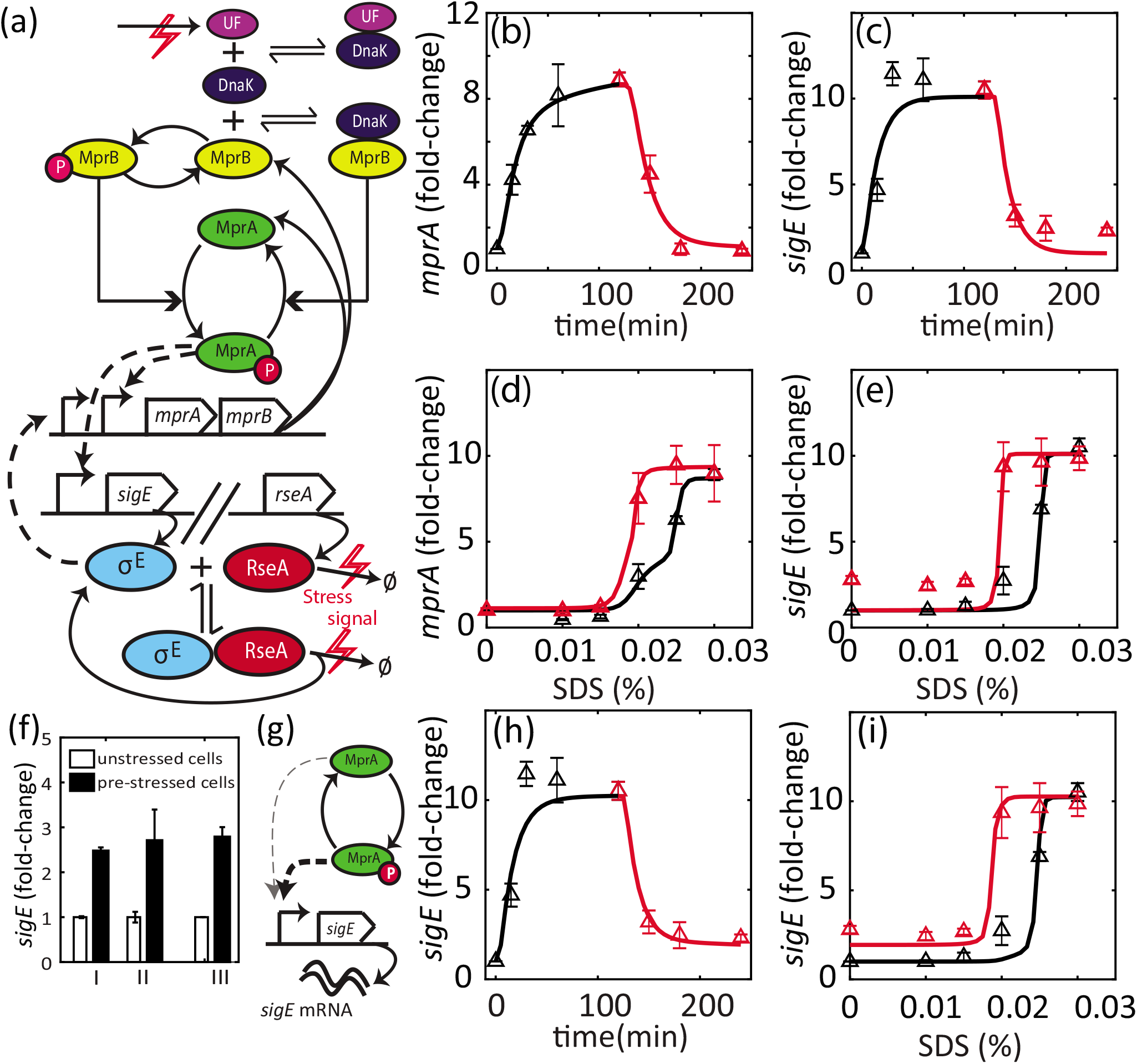
(a) Modification to MprA/B TCS: Chaperone DnaK binds MprB and suppresses autokinase activity. Processes affected by SDS exposure are denoted by red *↯* symbol. SDS leads to build up of unfolded/misfolded extra-cytoplasmic proteins (UF), which are targets of DnaK. DnaK subsequently releases MprB which is then free to autophosphorylate and catalyze phosphotransfer to MprA, thus activating the TCS. SDS exposure also leads to degradation of RseA (Ø indicates degradation products) and release of *σ^E^*. Dashed lines indicate transcriptional regulation. (b-d) Parameter fitting to model with modified TCS. This model can simulate close matches to time-course (b,c). Moreover, dose-response simulations show hysteresis depending on history of SDS exposure (d,e). However, residual *sigE* mRNA(e, red triangles) cannot be explained by this model. (f) Residual activity of *sigE* can be observed in multiple biological replicates. Measurement of *sigE* in the pre-stressed cells at 2hrs after removing SDS (solid bars) show nearly 2.5 fold more of *sigE* mRNA as compared to that in the unstressed cells (hollow bars). (g) Unphosphorylated MprA binds *sigE* promoter (with lower affinity compared to MprA-P). (h,i) Residual *sigE* activity can be explained by this hypothesis.

Given in-vitro studies showing that also unphosphorylated MprA can bind MprA-P target promoter DNA(7), we hypothesize that unphosphosphorylated MprA might act as a weak (i.e. with lower promoter affinity) transcriptional activator of *sigE* (Figure 4g). Addition of this interaction to the model (Methods) can explain this residual transcription activity, leading to a better fit overall (Figure 4 h,i). Thus the trade-off between hysteresis and response time is resolved. We illustrate this with a scatter of hysteresis and response-time for best-fitting parameter sets obtained using multiple particle swarm optimization runs (Figure S5). The points occupy the space at the intersection of experimental hysteresis and response time, left unoccupied by the previous model (Figure 3 f & Figure S5).

### Overexpressing DnaK from stress-responsive promoter partially abrogates stress-activation

If the hypothesized mechanism of hysteresis is accurate, we expect the dynamical properties of the network (activation level, hysteresis) to be strongly sensitive to DnaK concentration.The *dnaK* gene is essential for growth in *M. tuberculosis*, therefore deletion mutants are not viable (27, 28). Severe overexpression of DnaK has already been shown to abrogate induction of *sigE* and *mprA* mRNA following exposure to SDS (16). Here we test effects of DnaK overexpression on dose-response hysteresis. To perturb DnaK expression levels, we integrate into the genome an extra copy of *dnaK* expressed from the *mprA* promoter (Figure 5a; Methods). In this engineered strain, the native copy of *dnaK* expresses the chaperone constitutively, whereas the additional copy expresses it in a stress-dependent manner (Figure 5a). We argue that basal level of this promoter (in the absence of stress) may not be sufficient to fully attenuate stress-response.

**FIG 5.**
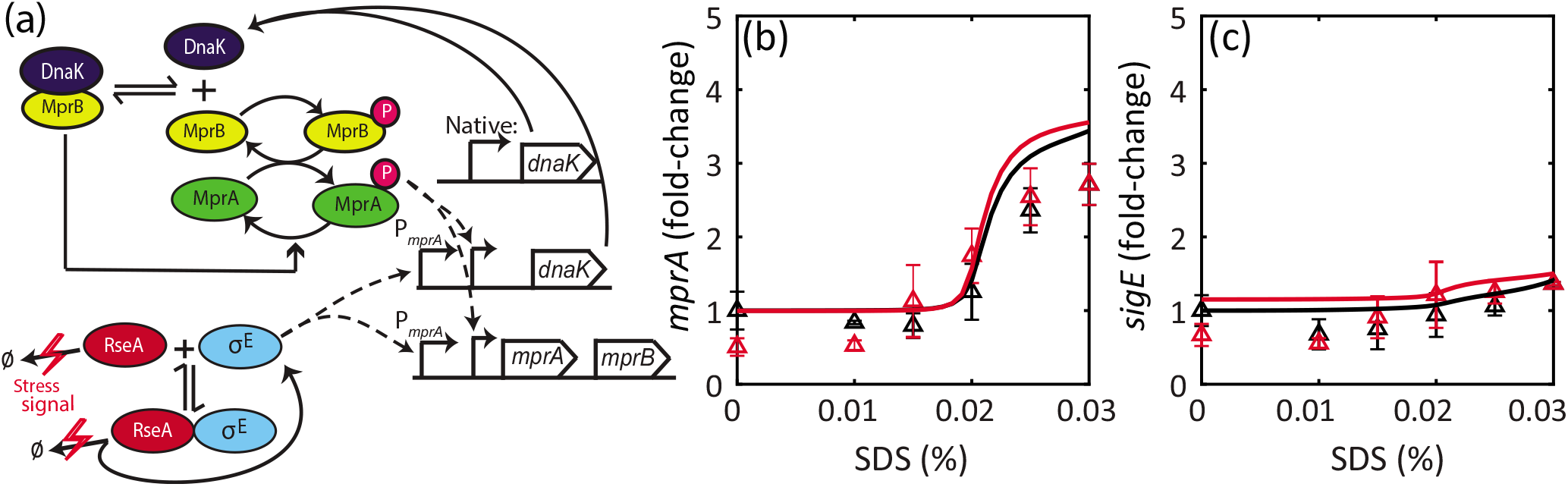
DnaK overexpressed from stress-responsive promoter (a) Schematic of MprAB-DnaK for the strain expressing DnaK from the *mprA* promoter (PmprA) in addition to the native copy. (b),(c) Much lower fold activation of *sigE* and *mprA* transcripts compared to wild type. DnaK-overexpressing cells were grown up to mid-log phase and treated with increasing SDS concentrations and harvested 2 hrs post-treatment (black triangles). DnaK-overexpressing cells were treated with bacteriostatic SDS (0.03%) for 2 hrs, centrifuged, and re-suspended in fresh media containing same or decreasing SDS concentrations and harvested 2 hrs post-treatment (red triangles). Solid lines show model simulations with DnaK translated from additional mRNA equal in concentration to mprAB mRNA (Methods).

With the engineered strain, we conduct dose-response experiments as described previously. We find that overexpressing DnaK results in attenuated activation of target genes (Figure 5 b,c). While the transcript levels of *mprA* increase nearly 3-fold over the range of SDS concentrations, *sigE* mRNA is not increased significantly. Since the *sigE* promoter is activated by MprA-P, this suggests that MprAB TCS is not activated following SDS treatment. On the other hand, modest upregulation of *mprA* could indicate activation of MprAB-*σ^E^* network through the alternative independent pathway of RseA degradation to release *σ^E^*. Our modeling results are consistent with this hypothesis. When we simulate the stress response of the engineered strain (Methods), it is possible to obtain qualitatively consistent transcript dose-response by tuning *dnaK* translation rate as a free parameter (Figure 5 b,c). Since *σ^E^*-RseA interaction in our model is completely independent of DnaK, we find that *σ^E^* activates the *mprA* promoter, leading to a modest upregulation. Induction of *mprA* is eliminated in the model in which no stress-dependent RseA degradation is present or if *σ^E^* is knocked out (Figure S6). In these simulations, DnaK is expressed at the basal level from the extra copy of *dnaK* that is nevertheless sufficient to inhibit MprB, which is also expressed at the basal level. Following SDS exposure, no significant upregulation of either is observed due to the loss of *σ^E^*-dependent activation of the *mprA* promoter driving the extra copy of *dnaK*.

We note that the experimentally observed change in stress response dynamics is not due to decrease in viability of engineered strain in SDS. Despite attenuation of stress response, at maximal SDS concentration used for hysteresis measurements (bacteriostatic concentration, 0.03%), cells remain viable over the experimental time-frame as shown by survival curves (Figure S7).

Taken together, these results suggest that increasing DnaK levels in *M. tuberculosis* leads to partial abrogation of stress response and suggest DnaK-dependent activation of MprAB TCS and DnaK-independent activation of *σ^E^*.

## DISCUSSION

The surface stress-responsive TCS MprAB, together with the alternative sigma factor *σ^E^*, forms a stress response network in *M. tuberculosis* that is a viable candidate for a bistable switch. Here we test the predicted bistable switch in the MprAB-*σ^E^* network in mycobacterial strains by measuring gene-expression in response to stress. Consistent with our previous prediction of bistability (15), we found that previously stressed cells show significantly higher levels of stress-activated transcripts compared to previously unstressed cells exposed to the same concentration of SDS. However, in contrast to predictions from our previous bistable model, we observed rapid accumulation of transcripts, suggesting that the assumed signaling network is inconsistent with experimentally observed dynamical properties. Our finding of a trade-off between hysteresis and response time in this model of MprAB-*σ^E^* network explains this inconsistency. We propose that the recently suggested mechanism(16) for activation of MprAB mediated by the chaperone DnaK can lead to bistability. Crucially, this mechanism does not result in a trade-off, and can explain all experimental observations. Furthermore, our model predictions of effects of DnaK perturbation are consistent with experimental measurements of engineered strains of *M. tuberculosis*.

We find that *M. smegmatis* neither displays hysteresis nor does it have a second positive feedback loop by which *σ^E^* regulates the *mprAB* operon. This results indicates that hysteresis is linked to mutual activation between MprAB and *σ^E^*. Since signaling architectures typically evolve in response to the requirement to survive in stressful environments (29, 30, 31) and that *M. smegmatis* is a non-pathogenic strain, it is tempting to speculate that the dynamical properties gained from the two positive feedback loop architecture in *M. tuberculosis* might be necessary for virulence(32, 33).

In modeling of two-component systems with a bifunctional kinase, it is commonly assumed that the activating singal simply increases autophosphorylation rate thereby decreasing the fraction of the unphosphorylated kinase that can act as a phosphatase. This assumption has been used in numerous modeling and theoretical analyses and has been sufficient to explain many observed dynamical properties of bacterial TCSs (23, 34, 35, 36). In our previous study predicting bistability in this network (15), autokinase rate of MprB was assumed to increase with stress. In contrast, in this study guided by the constraints set by our time-course and dose-response measurements, we uncovered that such activation assumption lead to a trade-off between hysteresis and response time. The trade-off is resolved by a more detailed activation mechanism involving the chaperone DnaK. Notably, the presence of a third protein stoichiometrically interacting the kinase will make systems response sensitive to changes in the concentrations of the two components. This is in contrast with absolute concentration robustness regime when third component is lacking(24, 25). It is interesting to see how potential loss of fitness due to lack of robustness in the DnaK-dependent activation mechanism may be compensated with fitness advantage due to a fast and sustained response of bistable network. Arguably the latter might be more important for virulent mycobacteria.

Involvement of chaperone DnaK with TCS signaling has precedents, since an effector protein regulating the activity of envelope-stress sensor kinase has been observed in other bacterial species. The response to envelope stress in *E. coli* is controlled by the CpxAR TCS. Athird component, CpxP, interacts with CpxAand suppresses its kinase activityin absence of stress (37). Upon conditions leading to overexpression of misfolded envelope proteins, CpxP is recruited away from CpxA, thus activating the TCS (38, 39). Overexpressing CpxP results in reduced Cpx response (38, 39). A similar mechanism exists in mammalian unfolded protein responses, where BiP acts as a folding chaperone for misfolded peptides exiting the endoplasmic reticulum. In addition to its role as a chaperone, BiP also binds and negatively regulates the activities of three transmembrane ER stress transducers: PERK, ATF6, and IRE1 (40). These three signaling proteins are released in conditions of increased load of misfolded peptides in the ER. Interestingly, overexpression of BiP leads to reduced activation of IRE1 and PERK. This could suggest a mechanism for detecting misfolded protein loads that allows for a rapid activation of stress response while sustaining activity even as the stress decreases. Therefore, our discovery of the dynamical consequences of a chaperone-mediated activation of signaling can help understand stress-response and protein homeostasis in diverse organisms.

## MATERIALS AND METHODS

### Experimental Methods

#### Bacterial strains, reagents, and growth conditions

*M. tuberculosis* H37Rv, *Mycobacterium smegmatis* (Mc2 155), and *Escherichia coli* XL1 blue (Agilent Technologies, Santa Clara, CA) were used. *M. tuberculosis* were grown in Middlebrook 7H9 broth (liquid) and Middlebrook 7H10 agar (solid) (Difco, Franklin Lakes, NJ), supplemented with 0.05% Tween 80, 0.2% glycerol, and 10% ADN (2% glucose, 5% bovine serum albumin, 0.15 M NaCl). However, 10% AND was excluded from the 7H9 or 7H10 media while M. smegmatis were grown. For DNA cloning, *Escherichia coli* XL1 blue (Agilent Technologies, Santa Clara, CA) were grown in Luria-Bertani (LB) broth or agar (Thermo Fisher Scientific, Waltham, MA). As needed, solid and liquid media were supplemented with 25 or 50 μg m/^−1^ kanamycin sulfate (Thermo Fisher Scientific, Waltham, MA) for Mycobacterium and *E. coli*, respectively. *M. smegmatis* knock outs in *mprA* (hygromycin marked) and *sigE* (zeomycin marked) were obtained from Prof. Thomas C Zhart (Medical college of Wisconsin) and Prof. Robert Husson (Boston Children Hospital, Harvard University), respectively, as kind gifts. *M. tuberculosis* knock-out in *mprA* was obtained from Prof. Issar Smith’s laboratory (PHRI center-Rutgers university), while a *sigE* mutant of *M. tuberculosis* were previously reported in Manganelli et al., 2001.

#### Construction of *P_mprA_-dnaK* fusion

DnaK was ectopically expressed from *mprA* promoter. For construction of *mprA* promoter::*dnaK* fusion, DNA fragments containing sequences 300 bp upstream of *mprA* plus first codon of the *mprA* open reading frame was PCR amplified and fused in frame with the N-terminal of *dnaK* coding region. Primers are listed in Table S1. The fusion construct was finally cloned into an integrative *E. coli*-mycobacteria shuttle vector pMV306-kan (Braunstein et al., 2001,Stover et al., 1991). Construction of the *P_mprA_-dnaK* fusion was verified by DNA sequencing. *P_mprA_-dnaK* fusion construct was electroporated in *M. tuberculosis* and the transformants were selected on kanamycin plates. Integration at the attp loci of the genome was verified by PCR.

#### SDS treatment

For gene expression analyses, mid-log cultures of *M. tuberculosis* and *M. smegmatis* were washed prewarmed (at 37°C) 7H9 medium, and treated with 0.03% and 0.02% SDS, respectively. These concentrations of detergent are bacteriostatic and had no cidal effect (Datta et al., 2015) (S Figure). Bacterial cultures were grown at 37 °C with shaking before and after SDS treatment. Gene expression analysis were performed from two sets of assays: SDS-time course with and SDS-concentration course. For time course experiments, mid-log cultures of *M. tuberculosis* and *M. smegmatis* were treated with specific bacteriostatic SDS concentration (mentioned above). 2 ml culture aliquots were harvested at various time intervals up to 2 hrs of SDS treatment, for RNA extraction. After 2 hrs of SDS treatment, part of the bacterial cultures was centrifuged and pellet was resuspended in SDS-free 7H9 broth (prewarmed) and incubated at 37 °C with shaking. Aliquots were collected at various time intervals up to 2 hrs of incubation in SDS free medium, for RNA extraction.

For SDS concentration course experiments, exponentially growing cultures were treated for 2 hrs with an increasing doses of SDS ranging from 0% to 0.03% (*M. tuberculosis*) or 0% to 0.02% (*M. smegmatis*). Aliquots were collected after 2 hrs of treatment for RNA extraction. After 2 hrs of incubation with highest SDS concentration (0.03% for *M. tuberculosis* or 0.02% for *M. smegmatis*), bacterial cultures were equally distributed in different tubes, centrifuged and pellets were resuspended in 7H9 medium (prewarmed) with decreasing SDS concentrations ranging from 0.03% to 0% (*M. tuberculosis*) or 0.02% to 0% (*M. smegmatis*). Bacterial cultures were further incubated at 37 °C with shaking for 2hrs. Aliquots were collected after 2 hrs for RNA extraction. As control, samples treated with same dose of SDS before and after centrifugation were tested to avoid any experimental artifacts.

#### RNA extraction and enumeration of bacterial transcripts

Details for RNA extraction and gene expression analysis was mentioned in Datta et al., 2015. Briefly, mycobacterial cells were disrupted by bead-beating (Mini-Beadbeater-16, BioSpec Products, Bartlesville, OK) in presence of 1 ml TRI reagent (Molecular Research Center, Cincinnati, OH). The aqueous phase was separated by adding 100 μl BCP Reagent (Molecular Research Center, Cincinnati, OH) and was collected after centrifugation.Total RNA was precipitated with isopropanol, washed with 75% ethanol, air dried and resuspended in diethyl pyrocarbonate (DEPC)-treated H2O for storage at −80 °C. Reverse transcription reactions were performed with ThermoScript™ Reverse Transcriptase (Invitrogen, Carlsbad, CA). Reverse transcription reactions were performed with random hexamer. Bacterial transcripts were quantitated by real time measurements using gene-specific primers and molecular beacons. Gene copy number was normalized to the 16s rRNA copy number (Shi et al., 2003). Nucleotide sequences of PCR primers and molecular beacons are listed in Table S2.

#### Survival analysis of wild-type and engineered *P_mprA_ – dnaK* strain

*M. tuberculosis* cultures were grown upto exponential phase using appropriately supplemented 7H9 liquid media. Exponentially growing cultures were treated with bacteriostatic (0.03%) and bactericidal (>0.03%) doses of SDS for 4 hrs or 24 hrs. Post treatment, bacterial cultures were incubated at 37 °C with shaking. Aliquots were plated for CFU enumeration before and after 4 hr or24 hrs of treatment. Input recovery was determined compared to CFU before the treatment.

### Computational Methods

#### MprAB-*σ^E^* network models

The dynamic ODE model describing MprAB, *σ^E^* and RseA concentrations was based on previous work from our team (15). We retained most of the model components, except for two changes. First instead of MprB autophosphorylation rate increasing with signal, we assume MprB existing in two conformations: kinase (MprB*) and phosphatase (MprB). We assume only the kinase form undergoes autophosphorylation, while only the phosphatase form can dephosphorylate MprA-P. The rate of conversion from MprB to MprB* (*k*_1_) was dependent on SDS concentration. Secondly, we introduced degradation of RseA (free or *σ^E^*-bound) with a rate (*k_rd_*) dependent on SDS concentration. Degradation of *σ^E^*-RseA results in a positive flux for *σ^E^*. Detailed transcription, translation and post-translational interaction reactions are described in Supplementary File 1. Instead of a first order activation of MprB, the DnaK-model includes a second order reaction, where MprB (kinase) binds DnaK into a complex (phosphatase).

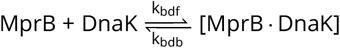

In presence of SDS, unfolded protein load (UF) builds up, leading to DnaK switching away from binding MprB to binding unfolded proteins:

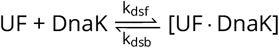

The total amount of UF remains constant and is dependent on SDS concentration. At t = 0, UF amount increases from 0 to UF(SDS). Upon deactivation of stress, we assume all UF is washed away, and all DnaK bound UF is freed during the washing. All reactions are summarized in Supplementary Material. Ordinary differential equations and parameter tables available in Appendix.

#### Signal-dependence

SDS concentration was incorporated into the models with a hill function. Parameters such as RseA degradation, phosphatase-to-kinase switching (canonical TCS model) and DnaK target concentration (DnaK model) were dependent on SDS with a saturating function as shown below:

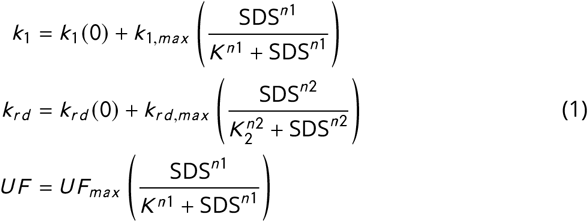

### Time course and dose-response simulations

Input signal enters a model at two points: MprAB activation (different depending on the model) and *σ^E^* activation (through RseA degradation).

#### Time course simulation

Steady state of the ODE model is initially simulated at no stress. At t=0, both signal parameters increase to the level corresponding to high SDS (0.03%) concentration. With pre-stress steady state as initial condition, the ODEs are simulated to obtain a time course of *mprA* and *sigE* mRNA between t=0 and 120 min (see supplementary text). The values are normalized to respective pre-stress levels to obtain a fold change mRNA time course for stress activation. Using state at t=120min is used as initial condition and setting signal parameters corresponding to 0 SDS, the deactivation time course of mRNA is obtained.

#### Dose-response simulation

The above pre-stress initial condition is also used to numerically compute transcript levels at t = 120 min, for parameters corresponding to each intermediate SDS concentration (Figure 3c, d). This gives us the simulated dose-response for unstressed cells. The state at t = 120 min at high SDS is used as initial condition to compute transcript levels at same intermediate concentrations to give the simulated dose-response for previously stressed cells. All ODE solutions were obtained with the ode15s solver in MATLAB.

### Error calculation and parameter fitting

Experimental mRNA was measured using qRT-PCR and normalized to 16S rRNA as internal control. Each mRNA measurement (at different time points, or different treatment conditions) was then normalized to the unstressed measure to generate a fold change value. Simulated mRNA levels were also normalized to unstressed levels. Error was then calculated as a sum of squared residual of corresponding simulated and measured fold change values at the last time-point *t_f_* = 120 min.

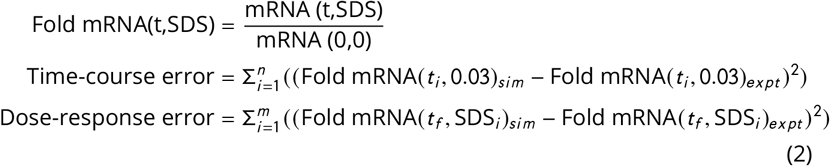

Parameter fitting was performed using particle swarm optimization with MATLAB, using the above error as objective function. Many of the kinetic rate constants have not been measured experimentally in *M. tuberculosis*. Given that the number of data points is low, and the number of parameters is high, a family of parameter sets was obtained for each model to account for “sloppiness” in parameters.

#### Analysis of simulations

Response time (τ_90_) was estimated as the time point afterstimulus at which simulated mRNA level was 90% of that at steady state. For experimental data, τ_90_ was estimated as the window between the latest data point at which the mRNA level is below 90% and first data point at which it is above 90% of the final value (at 2hrs post stress). Degree of hysteresis was calculated as a mean of difference between fold mRNA levels for previously stressed and not stressed cells (red and black markers respectively in Figure 3c,d) at two intermediate SDS levels (0.02% and 0.025%).

## SUPPLEMENTAL MATERIAL

**FIG S1.** Survival of wild-type *M. smegmatis* at increasing SDS concentrations. Colony forming units (CFUs) increase at 0-0.01% SDS, remain relatively constant at 0.02% SDS and decrease at 0.03% SDS.

**FIG S2.** ODE model of MprAB-*σ^E^* shown in Figure 3 e is analyzed using time-scale separated modules as discussed previously in ref (15). Both types of parameter sets (Figure 3 a-d) used to obtain transcription module (red) and post-translational module (blue) at 0.02% SDS. Parameter sets of type II show bistability, as is seen by the 3 intersection points for blue and red curves (solid). Parameter sets of type I (dashed) do not show bistability. Intersection occurs in the range of saturation where MprA-P is insensitive to changes in total MprA. Thus there can only be one intersection point.

**FIG S3.** To investigate the presence of bistability of the ODE model for *M. tuberculosis* MprAB-*σ^E^* network with DnaK, we compute a bifurcation diagram forthe parameter SDS (blue). The bifurcation diagram shows a range of SDS values where the model is bistable. Similar bistability range is observed if the steady state is found by integration of the ODE model starting from uninduced (steady state of SDS=0, black line) or fully induced (steady state of SDS=0.03%, red line) initial conditions.

**FIG S4.** MprAB-*σ^E^* stress-response network model can be adapted for *M. smegmatis. M. smegmatis* lacks feedback from *σ^E^* to *mprA*. We adapt the *M. tuberculosis* model to *M. smegmatis* by removing this feedback loop and altering protein and mRNA degradation rates. We assume that activation mechanism of MprAB as well as *σ^E^* is conserved between the species. We then fit the model to *M. smegmatis* data (time course shown in a, b as well as dose response shown in c, black and red squares). The model can explain time course data while not displaying dose-response hysteresis (overlapping red and black lines, c)

**FIG S5.** Analysis of multiple parameter sets that generate fits of the type in Figure 4. (a) Magenta circles represent best-fitting parameter sets for MprAB-DnaK model as in Figure 4a. (b, c) Simulations with unphosphorylated MprA as a weak activator match time course in addition to dose-response shown in Figure 4h, i

**FIG S6.** Simulations of the model with an extra copy of *dnaK* expressed from the *mprA* promoter performed with *in silico sigE* deletion mutant (top row, a,b) or with no stress-dependent increase in RseA degradation (bottom row, c,d) show no upregulation of *mprA* (in contrast to Fig. 5b)

**FIG S7.** Survival of wild-type and engineered DnaK strain shown in Figure 5. Recovery of colony forming units (CFUs) after 4 hrs of exposure to SDS at the concentrations bacteriostatic (0.03%) or bactericidal (**>**0.03%) for wild-type strain(18). Survival of wild-type and DnaK overexpressing strain was comparable when treated with bacteriostatic dose of SDS (0.03%). However, treatment with bactericidal doses of SDS showed that DnaK overexpression strain was more susceptible to killing compared to wild-type strain.

**TEXT S1.** Model reactions, ODEs and parameters used in this study.

## ACKNOWLEDGMENTS

The research was supported by Welch Foundation Grant C-1995 to OAI, NIH grants GM 096189 to OAI (co-PI: MLG) and AI 122309, AI 104615, and HL 149450 to MLG. We thank Angela Cintolesi for her contributions to the mathematical models and parameter fitting algorithms.

## Notes

### Competing Interest Statement

The authors have declared no competing interest.

